# Deciphering Coccolith Formation: Advanced Microscopy Insights from the Biomineralisation of *Gephyrocapsa huxleyi*

**DOI:** 10.64898/2026.04.08.717164

**Authors:** Alexander Triccas, Mariana Verezhak, Johannes Ihli, Manuel Guizar-Sicairos, Mirko Holler, Fraser Laidlaw, Martin R. Singleton, Virginie Chamard, Rachel Wood, Tilman Grünewald, Fabio Nudelman

## Abstract

Coccolithophores are unicellular marine phytoplankton that produce complex and intricately shaped mineralised scales called coccoliths. Coccoliths are produced in an intracellular vesicle where crystal nucleation occurs, from which several individual calcite units develop with anisotropic crystallographic facets, prompting studies into the cellular mechanisms which control crystal growth within the cell. Here, we characterise those morphological developments in 3D that occur during the formation of coccoliths by the species *Gephyrocapsa huxleyi* using cryo-ptychographic X-ray computed tomography. This technique is ideally suited to study coccolith mineral development, as intracellular structures can be imaged intact in their native state without needing to disrupt cells. Combined with additional imaging of developing coccoliths using cryo-transmission electron microscopy and scanning electron microscopy, we report the developmental stages involved in coccolith growth across the complete mineralisation period, while also showing that the constrained space created by individual crystal units growing in close confinement affects the final crystal morphology and overall mineral structure. These findings provide clarification on the mineralisation pathways that coccolithophores and other biomineralising organisms use to control the formation of highly functionalised crystalline structures, particularly relevant in the design of materials with tunable properties.

## 1. Introduction

Biomineralised tissues are essential to the survival of organisms across all six kingdoms, with their formation precisely controlled by manipulating the fundamental pathways of crystallization. By employing an array of cellular mechanisms, numerous aspects of crystal formation are modified: nucleation and growth stages, as well as morphology and polymorph selection [1-4]. As a result, complex-shaped crystals with hierarchical organisation form, with vastly different structures than their synthetically-grown counterparts. The ability organisms have to produce a range of functional mineralised structures with tunable properties from readily available ions attracts significant interest in materials science [5].

A model organism in the study of biomineral formation are unicellular marine phytoplankton called coccolithophores. These cells produce remarkable micrometer-sized scales with complex morphologies called coccoliths, displaying multi-faceted intricately shaped crystals that form under strict biological regulation. Coccoliths consist of several interlocking single calcite subunits, synthesized in an intracellular vesicle where crystal nucleation and growth take place [6, 7]. When fully mature, coccoliths are exocytosed from the cell, creating an external layer of interlocking scales called the coccosphere [8]. While coccolith formation is a well-studied process, many outstanding questions remain, particularly as to the conditions under which nucleation and growth are controlled in the intracellular vesicle. Crystallisation pathways must also be resolved to determine the energetic costs involved in coccolith formation, which will largely affect how mineralisation will respond to an increase in ocean temperature and acidity in light of anthropomorphic climate changes [9-12]. This is of significant importance given the role that coccolith mineralisation has in the global carbon cycle through its contribution to the formation of large geological carbon reservoirs [13].

Numerous studies have contributed to our understanding of how crystal growth and morphology is controlled in coccolithophores by defining the growth stages required to produce coccolith mineral. Initially, coccolith formation was studied by extracting undeveloped mineral units from cells and characterising the crystal morphology at various stages in the V/R model [14-17]. Rhombohedral crystals first nucleate with their crystallographic *c*-axis orientated either vertically (V-units) or radially (R-units) aligned to an organic base plate [18, 19]. Individual units then develop through mineral addition, with the expression of several specific crystallographic facets indicating that growth is precisely controlled. Several cellular mechanisms have been proposed to influence biological crystal formation in coccolithophores. Organic macromolecules are often cited as a major control factor of crystal growth. Polysaccharides extracted from mature coccoliths have been shown to promote anisotropic growth of specific crystallographic facets in synthetically produced calcite [20, 21], while lattice distortions within the coccolith mineral indicate that macromolecules could template growth by blocking certain facets [22]. Growth is also known to be controlled through physical constraints created by the close confinement of coccolith subunits, with crystallographic facets dictated by the prevention of mineral expansion by nearby structures [23, 24]. Additional control is imparted through the intracellular vesicle where mineralisation occurs, both through physical confinement and alteration of ionic concentration gradients [25, 26]. A feature of coccolithophore biomineralisation is that each of the 200-250 classified living species produce coccoliths with completely unique organisation and morphology of their crystal subunits [12]. It is likely that each coccolithophore species employs these growth control mechanisms to differing extents, reflected in their structural and morphological diversity.

Difficulties in defining these species-specific control mechanisms comes from an incomplete understanding of how crystal morphology changes over the course of coccolith formation, particularly at early stages of growth. By focusing on the first stages of coccolith development in the species *Gephyrocapsa oceanica*, it was revealed that the vertical tube element is actually composed of two crystal layers contrary to the one in the V/R model, tilted in opposite non-vertical angles relative to the base plate [23]. This adds complexity to the overall crystal morphology reported for this species and is an important feature which allows units to become tightly interlocked and influences the later formation of the shield elements. It is possible that these features develop due to the highly constrained mineral growth space at these early stages, but this is difficult to clarify as undeveloped crystals are challenging to image in their native state.

Further characterisation of coccolith morphology across the entire window of mineralisation is therefore required to accurately determine the mechanisms involved in controlling crystal growth. To achieve this, entire cells of the coccolithophore *Gephyrocapsa huxleyi* (formerly known as *Emiliania huxleyi*) were imaged by cryo-ptychographic X-ray computed tomography (cryo-PXCT), which preserves cellular material in its native state and enables the 3D reconstruction of coccoliths in their entirety [27, 28]. As the field-of-view is sufficient to image around 80 *G. huxleyi* cells in an actively calcifying state, we were able to visualise a large number of coccoliths at multiple stages across their development and through 3D structural reconstructions can report quasi-continuous growth across the entirety of mineralisation. Additional characterisation of crystal morphology was provided by cryo-transmission electron microscopy (cryoTEM) and scanning electron microscopy (SEM). In sum, this allows a precise depiction of the sequence of coccolith growth, as well as reporting new findings on the influence that confined growth space has on crystal morphology and tight cellular control over the location of mineralisation. Furthermore, we provide an estimation of the mineral mass of individual coccoliths, both fully formed and at different stages of their maturation. These findings contribute to an overall aim of clarifying the cellular mechanisms that coccolithophores and other biomineralizing organisms employ to control crystal growth in the design of hierarchical functional structures.

## 2. Experimental

### 2.1 Cultures

Cultures of *Gephyrocapsa huxleyi* (920/9) were obtained from the Culture Collection of Algae and Protozoa (CCAP), Oban, Scotland. Cultures were incubated at 18 °C on a 12:12 light-dark cycle. *G. huxleyi* was grown in K/2 medium, adapted from Keller et al., 1987 [29], prepared using natural seawater from St Abbs Marine Station. Cultures were also supplemented with a Penicillin/Streptomycin/Neomycin Antibiotics solution (Fisher BioReagents). Cultures were maintained by subculturing 1 mL of culture into 30 mL fresh medium every 2 weeks. Cell growth was measured using a Leica optical microscope and Neubauer improved haemocytometer.

### 2.2 Cryo-ptychographic X-ray computed tomography (cryo-PXCT)

#### Preparation of G. huxleyi cells

Cells were prepared by decalcifying 5 mL of culture in its exponential phase (5 days after subculturing) with 0.1 M pH 8.0 EDTA to dissolve external coccoliths. Cells were left overnight in low-calcium (100 µM) Aquil artificial seawater, preventing further coccolith growth in order to recalibrate calcification. A 0.1 M CaCl_2_.2H_2_O (100 µL) solution was then added to raise [Ca^2+^] back to seawater levels (10 mM) and left for 24 h to provide sufficient time for enough coccoliths to form to fill the entire coccosphere. Cells were harvested in 1 mL aliquots, centrifuged at 1500 G, and concentrated to a ∼50 µL volume.

Capillaries with tips 10 µm wide were produced using a Flaming-Brown capillary puller (P-2000, Sutter Instruments) and mounted in hollow OMNY pins with UV resin [30]. The samples were loaded into the capillary tips from the accessible backside. The samples were then further sedimented into the very tip of the capillary by a small settling period or by mild centrifugation (2000 G for 240s). The capillaries were rapidly plunge-frozen in liquid ethane using a vitrification robot (FEI Vitrobot Mark VI). The sample chamber was maintained at 21 °C and 100% humidity. Particular care was taken to minimize the time lag between sample preparation and sample freezing. The samples were stored at cryogenic temperature under liquid nitrogen and transferred to the beamline using a transport dewar to ensure a continuous cryogenic cold-chain, which was kept up until after the samples were measured in the OMNY instrument [31].

#### Tomogram Acquisition and Reconstruction

Cryo-PCXT measurements were carried out at the cSAXS beamline (X12SA), Swiss Light Source, Paul Scherrer Institute, Switzerland with the OMNY instrument [31]. Ptychographic scans were performed at a photon energy of 6.2 keV under cryogenic conditions at 90 K. Doing the measurements under cryogenic conditions was important both for the preservation of the cellular material in its close to native state, and to minimize radiation damage that is inherent to the large dose needed to produce a complete dataset. An Eiger 1.5M detector, placed 7.2 m downstream of the sample was used to collect far-field intensity patterns at each scanning point. The same ptychography projection scan was performed at various equiangular orientations, rotating the sample about the vertical axis in the range between 0 and 180°. The exact imaging parameters are listed in Supp. Table 1.

The ptychographic reconstructions were carried out using 300 iterations of the difference-map algorithm, followed by 300 iterations of maximum-likelihood refinement [32], resulting in a reconstructed pixel size of 32 nm (Supp. Fig. 1) [33]. The projections were further processed and aligned using in-house developed matlab scripts (which are publicly available) [34]. Fourier Shell Correlation was used to provide a first estimate of the spatial resolution. The datasets were split angularly into two separately reconstructed datasets and compared using the Fourier shell correlation (FSC) approach [35]. In this way, the half-bit resolution was estimated as ∼56 nm. The corresponding FSC curve is presented in Supp. Fig. 1.

### 2.3 Cryo-Transmission electron microscopy (cryoTEM)

To extract intracellular coccoliths from *G. huxleyi* cells, 50 mL cell cultures early in the exponential phase (2 days after subculturing) were decalcified using 0.1 M pH 8.0 EDTA. The seawater medium was aspirated, and the pellet of decalcified cells was placed in a 0.1 M Na_2_CO_3_ solution (40 mL) overnight to cause lysis by osmotic pressure. The solution was concentrated down to a 20 µL volume and an aliquot (3 µL) of this coccolith-containing solution was applied to a cryoTEM grid and plunge frozen in liquid ethane at liquid nitrogen temperatures using a vitrification robot (FEI Vitrobot Mark VI). The sample chamber was maintained at 21 °C and 100% humidity. CryoTEM (Au/C, R2/2 Quantifoil Micro Tools Gmbh) were plasma treated for 1 min prior to freezing using a Quorumtech Glow Discharge system. An FEI F20 Tecnai electron microscopy with 200 keV field emission gun, equipped with a Gatan cryoholder operator at *ca*. -170 °C was used for imaging. Images were recorded on a Gatan K2 Summit camera.

### 2.4 Scanning electron microscopy (SEM)

For SEM imaging of mature *G. huxleyi* coccoliths and cells, 200 µL of culture in its exponential phase (5 days) was deposited on Isopore 0.2 µm PC membranes (Merck Millipore Ltd.). Cells were washed with 1 mL 8.0 pH dH_2_O to remove salt from the membrane and air-dried overnight. The membrane was mounted on SEM stubs and coated with Pt. For SEM imaging of intracellular coccoliths, cells were subjected to osmotic pressure using a 0.1 M Na_2_CO_3_ solution as described above for cryoTEM. A 200 µL aliquot of a concentrated coccolith solution was then deposited onto a 0.22 µm PC membrane. All cells and coccoliths were washed with 3 mL dH_2_O adjusted to pH 8.0 to remove salt from the membrane without dissolution of the mineral structures. All samples were coated with a thin layer of carbon before imaging. SEM imaging was performed on a Zeiss Crossbeam 550 cryoFIB-SEM with an accelerating voltage of 3 kV.

## 3. Results

### 3.1 Mapping morphological changes across coccolith development using ptychographic computed X-ray nanotomography

To ensure cells were actively calcifying when cryogenically preserved, mineralisation was induced by dissolving external coccoliths and then placing cells back in normal culturing conditions for 24 h. The resulting cells were directly frozen in solution and imaged by cryo-PXCT, which enables mineral structures and other dense features to be visualised and reconstructed in 3D (Supp. Fig 2a-c and movie S1). From 2D slices through a representative cell (Fig. 1a-f), a mineralising calcite coccolith (red arrows in Fig. 1a-d) can be observed within, while several coccoliths (yellow arrows in Fig. 1a-f) surround the outer surface. 3D volume renderings of these features (Fig. 1g-h) reveal the intracellular coccolith (red Fig. 1g-h) is not fully developed, in comparison to the extracellular coccoliths (yellow Fig. 1h) that cover the cell surface. The morphology of the mineralised cell (Fig. 1i) appears to be consistent with those imaged by SEM (Fig. 1j), suggesting structural reconstructions from cryo-PXCT are accurate and that minimal damage occurred from cryogenic preparation. Electron-dense bodies can also be observed within cells (black arrows in Fig. 1f and black in Fig. 1g-h), similar in size and morphology to ion-rich bodies defined in this species [36]. Cryo-PXCT was used to report the variable morphology of similar bodies related to Ca storage and transport in the coccolithophore *Chrysotila carterae* [37], however we could not observe dense bodies with any consistent structural features or spatial relationship to the intracellular mineral structures in this study.

**Figure 1.**
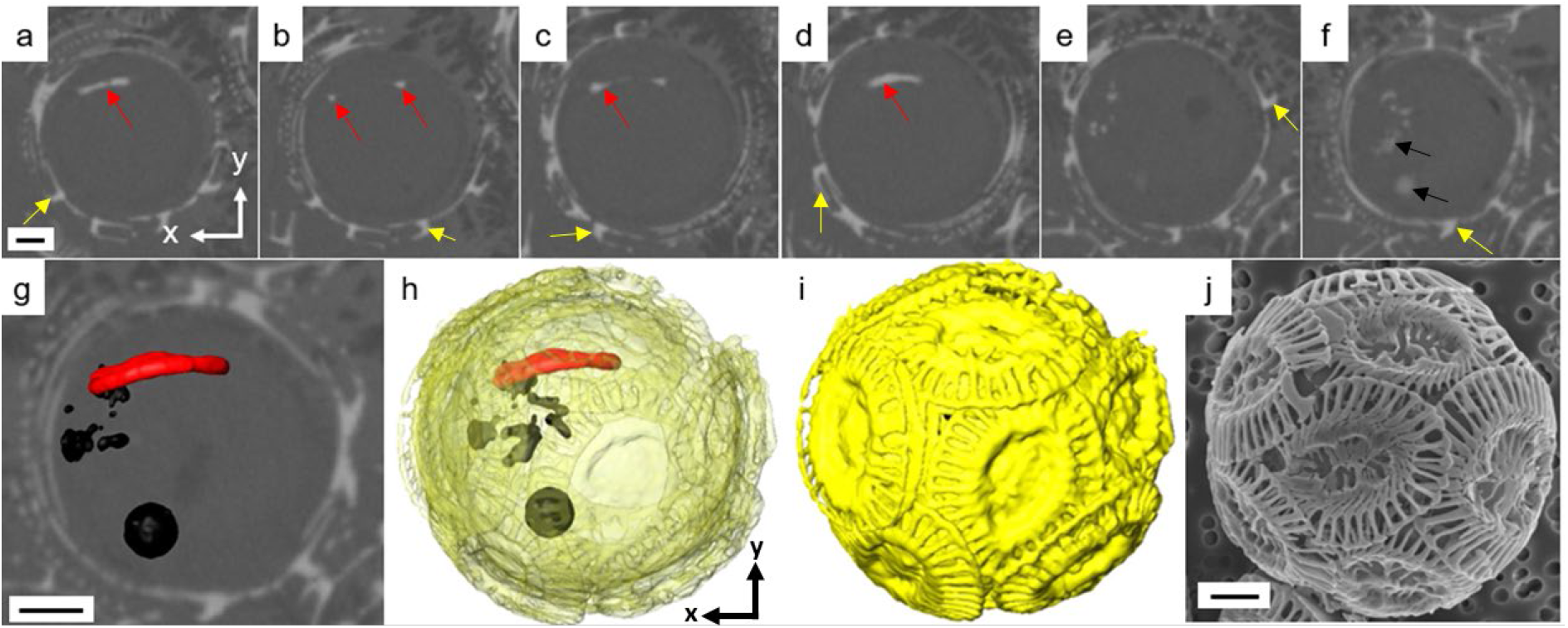
cryo-PXCT of an actively calcifying *G. huxleyi* cell. Shown in (a-f) are orthoslices through one calcifying *G. huxleyi* cell, highlighting forming intracellular coccolith (red arrows) and electron-dense bodies (black arrows), alongside several interlocked extracellular coccoliths (yellow arrows) on the outer cell surface. Provided in (g) are volume renderings of the intracellular coccolith (red) and electron-dense bodies (black) displayed in the ortholsices in (a-f). (h) Outside the cell, external coccoliths (yellow) cover the surface. The full coccosphere reconstruction shown in (i), in comparison with a scanning electron micrograph of an intact cell (j). Scale bars: 0.5 µm.

From the population of *G. huxleyi* cells inside the capillary (Supp. Fig. 2a-c), a total of 88 intracellular coccoliths were identified. These coccoliths had distinct morphologies related to the stage of growth they were in when cryogenically preserved. Presented in Fig. 2 is a series of cross-sections and volume renderings of coccoliths selected to highlight the morphological transformations occurring across mineral formation, which are categorised here into five development stages based on clearly identifiable structural features and the assumption that a coccolith can only gain weight and volume during its growth. The smallest mineral structures which constitute Stage 1 were flat spherical rings, possibly related to the first nucleation of the protococcolith ring. The smallest rings had a diameter as low as ca. 180 nm (Stage 1a), but most were thicker, with a diameter between 250-300 nm (Stages 1b). Mineral growth appears to be isotropic at this early stage. Around half the intracellular coccoliths we observed were at growth stage 1, with others distributed evenly across the later stages (Supp. Table 1). This suggests that this stage may contain the rate limiting step in coccolith formation.

**Figure 2:**
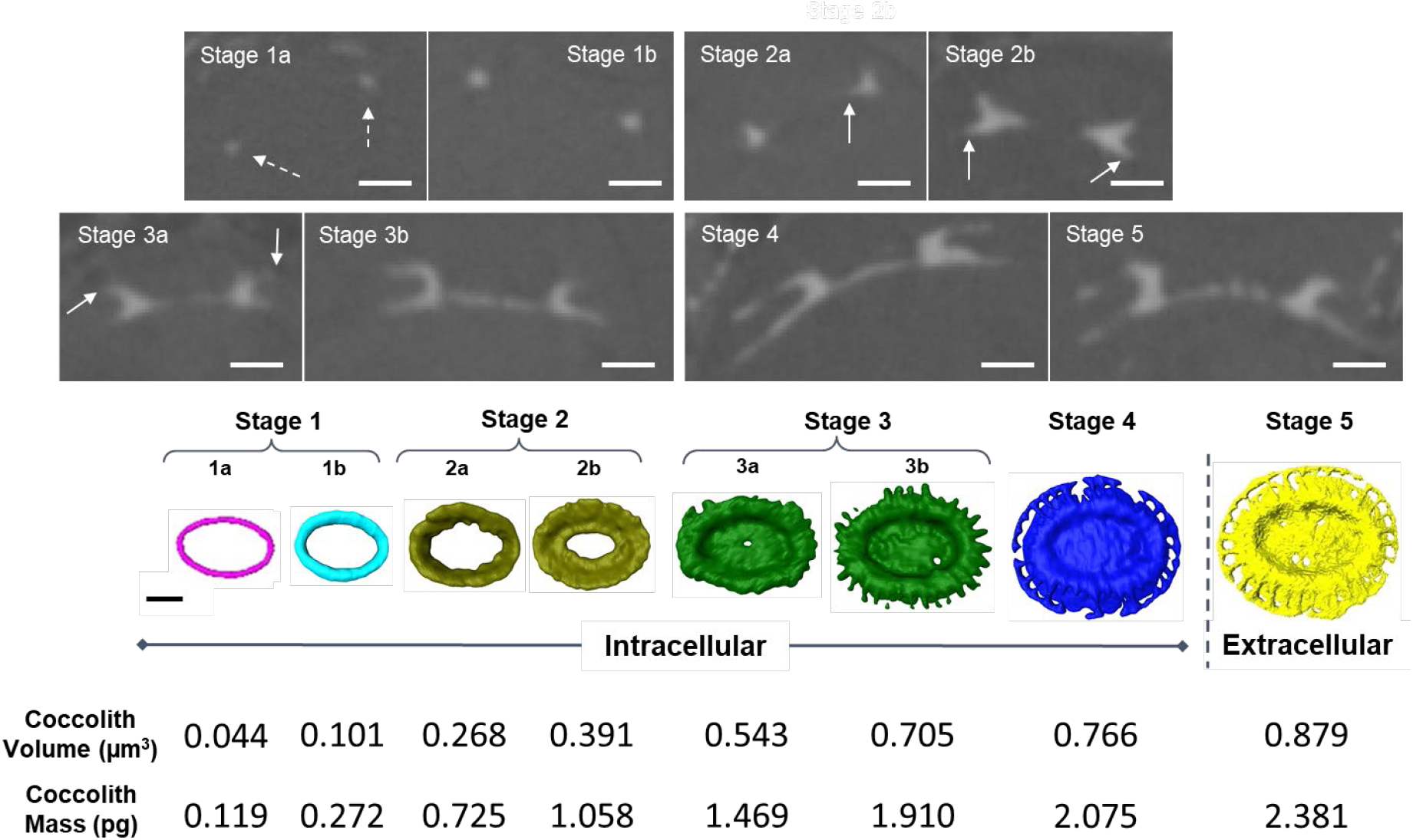
Morphological development of *G. huxleyi* coccoliths. Top: Cryo-PXCT provides slices through the 3D volume of coccoliths at various stages of development, showing morphological changes as the mineral becomes fully formed. Arrows indicate first appearance of: thin mineral ring in stage 1, central area elements in stage 2a, proximal shield in stage 2b, distal shield in stage 3a. Bottom: Volume rendering of the 8 coccoliths, with the calculated volume and mass of coccoliths at each stage of development. Scale bars: 0.5 µm.

Stage 2 is defined by preferential mineral addition to the top side of the spherical ring, relating to the vertical crystal growth that causes the tube element to form. Initial vertical growth results in a coccolith at Stage 2a with a height of 300 nm. Additionally, small extensions also appear from the base of the inward facing side of the mineral (arrows in stage 2a, Fig. 2), indicating the first appearance of the central area elements. Further vertical growth in Stage 2b leads to the completion of the tube element, with a height of 400 nm. This stage also coincides with the coccolith tube expanding at its base, with outward addition starting the formation of the proximal shield (arrows in stage 2b, Fig. 2). In combination with this, larger mineral protrusions also appear on the inwards facing side of the coccolith ring. These additions visibly change the coccolith morphology, which is now 500 nm wide at its base.

Stage 3 is defined by the presence of mineral regions which connect across the base of the coccolith tube, representing the completion of central area elements. Further identification of coccoliths at this growth stage was distinguished by new mineral regions at the top of the tube element (arrows in stage 3a, Fig 2). Rather than continuing to extend vertically, the mineral curves around as growth changes to an outward direction, reflecting the formation of the distal shield elements. Both the proximal and distal shield elements then grow outwards in the same radial direction (Stage 3b), both extending by approximately 200 nm from the coccolith tube.

Stage 4 is classified as fully developed coccoliths, with T-shaped distal shield elements. In a mature coccolith, the distal and proximal shield elements are respectively 680 and 800 nm in length. The thickness varied across both units, ranging between 30 to 120 nm. It should also be noted that contrary to previous reports of single coccoliths being produced at one time, we observed multiple structures forming within the same cell. In all cases, one coccolith was always fully mature, while the other was either early in development around Stage 1-2 (Supp Fig. 3a) or nearing full maturation (Supp Fig. 3b). Stage 5 is morphologically identical to Stage 4, with coccoliths located outside the cell rather than inside, reflecting exocytosis of the scale rather than mineral growth.

PXCT also offers the advantage of providing electron-density values which relate to composition of the structure imaged. Electron density can be converted into volume and mass, and as cells are imaged intact, the size and weight of coccoliths throughout their growth can be estimated. To calculate these values in light of the effect that resolution limitation has on identifying voxels belonging to the outer coccolith mineral regions, electron density values of all voxels comprising the coccoliths were summed together and divided by the expected electron density value for calcite mineral (0.82 n_e_Å^-3^) [27]. This value was then multiplied by the size of a single voxel (3.28 × 10^-17^ cm^3^) to obtain the volume of the whole structure. From the volume of the coccolith, the mass can be calculated by multiplying by the calcite mass density (2.71 g/cm^3^) [38, 39], both displayed alongside coccoliths in Figure 2. The mass of all coccoliths analysed is shown in Supp. Fig. 4.

While coccolith volume and mass generally increased as the structure became more developed, substantial variability in values across the same growth stage causes some coccoliths at later stages of development to have lower masses than those at earlier stages, particularly evident across stages 2-3 and 3-4. In some cases, this can be explained by malformations in the coccolith, particularly at stage 4, but when no visible gaps in the structure are present, these discrepancies likely represent biological variabilities in the size of the scales [40]. The calculated volume and mass of the mature coccolith displayed in Fig. 2 was consistent with literature values calculated from X-ray nanotomography [40] and profiles estimated from SEM images [41]. We additionally observed mature intracellular coccoliths with a range of masses (Supp. Fig. 4), once again highlighting the biological variability in the size and mass during coccolith production.

By reporting changes of masses at different growth stages, we are able to provide rough estimates for the proportion of mineral in each element of the coccolith. The mineral mass of the smallest structure observed was 0.087 pg, although the rest of the coccoliths across growth stage 1 had masses ranging between 0.119-0.436 pg. This suggests that during the initial radial expansion of the tube element, the mineral mass of the structure increases two-to three-fold. As soon as vertical growth begins at stage 2a, the mass added to the coccolith increases substantially, reaching over 1 pg, highlighting that the highly mineralised tube element contributes nearly half of the estimated coccolith mass. The mass increase observed during Stage 3a reaching up to 1.469 pg, most likely comes from the central area elements. The last stages (3b to 4) involve the full formation of shield elements, coccoliths reaching between 2 and 3.6 pg in mass. This highlights the considerable proportion of mineral mass contained within relatively thin distal and proximal shield elements, around a third of the total coccolith weight.

Using cryo-PXCT we also mapped the 3D position of each coccolith within the cell. This revealed that coccolith location appears to be strongly dependent on its stage of growth (Fig. 3a-e). At the beginning of their development (stage 1), coccoliths were mostly located in a central region of the cell (red, Fig. 3a-b). As the coccolith developed across Stage 2, scales were generally closer to the cell periphery (red, Fig. 3c), and by Stages 3 and 4 coccoliths were always located with their convex side at least 50 nm away from the plasma membrane (red, Fig. 3d-f). Additionally, mature coccoliths were always located in regions of the cell that were not covered by external coccoliths (yellow points, Figs. **3d-e**), even when only small gaps between two non-interlocking scales were available when the coccosphere was nearly complete (yellow points, Fig. **3f**). This suggests that the location where exocytosis takes place is carefully controlled by the cell. Although we did not view a coccolith being exocytosed, scales are removed with their convex side facing the plasma membrane, contrasting other coccolithophores species which expel scales in the opposite direction before they undergo extracellular rotation in order to interlock with other outward facing scales [8].

**Figure 3:**
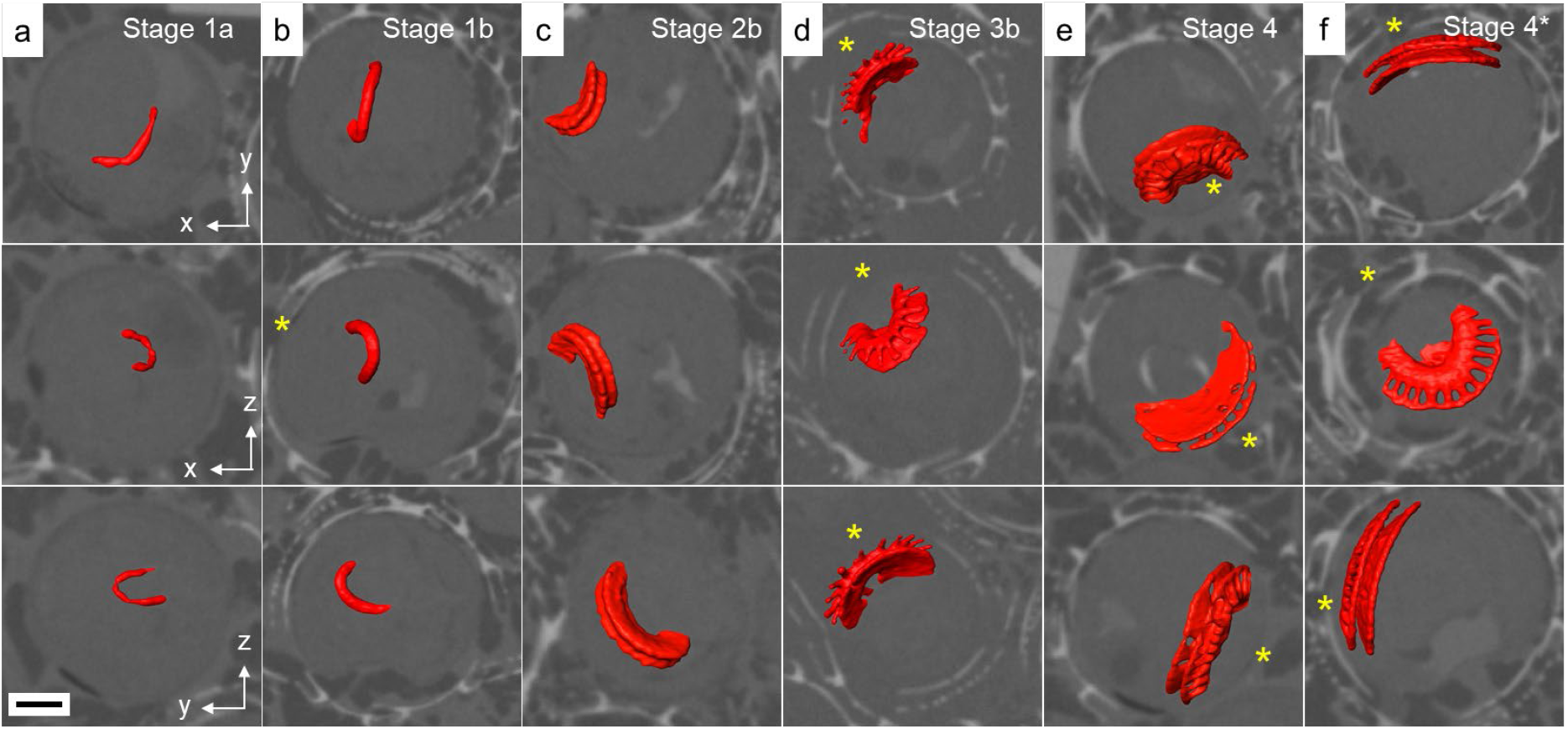
Cryo-PXCT tomograms showing cellular position of intracellular *G. huxleyi* coccoliths. (**a-e**) Cross-sections through the volume of 6 separate *G. huxleyi* cells containing coccoliths at different stages of development. Volume renderings of forming coccoliths are shown in red. In stage 1 (**a-b**), coccoliths are found in the centre of the cell. Between stages 2-4 (**c-f**), coccoliths are found in proximity to the membrane. Yellow markers indicate gaps in the coccosphere. Scale bar: 0.5 µm.

### 3.2 Changes in crystal morphology across coccolith mineralisation

While cryo-PXCT enabled changes in morphology across the majority of coccolith mineralisation to be identified, the individual crystals that comprise the structure could not be resolved by the technique. To fill in these gaps and provide information of how crystal morphology develops during mineralisation, intracellular coccoliths were extracted from calcifying cells and imaged using cryoTEM and SEM. We could identify coccoliths at various growth stages, including the earliest formed protococcolith ring which is composed of rhombohedral crystal units (Fig. 4a-c). Both R-units and smaller V-units (white arrows Fig. 4b) were visible. This could be representative of a coccolith at growth Stage 1a, although crystal units were not directly in contact with each other but rather appear to be connected by a layer of material (arrows, Fig. **4c**) which could be the base plate rim where crystals nucleate.

**Figure 4:**
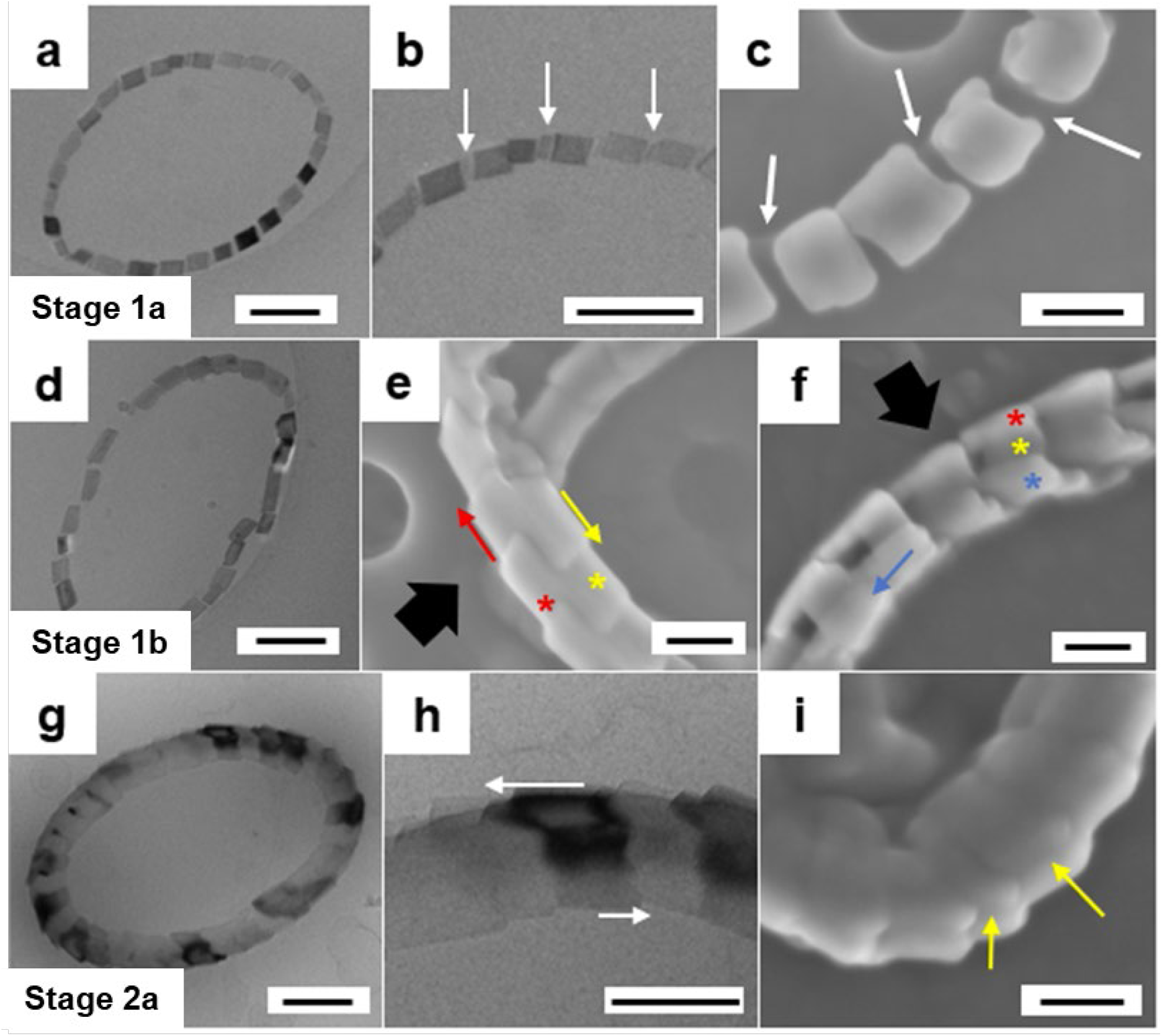
Formation of the outer and inner tube elements through Stages 1 and 2a. (**a**) cryoTEM image of a coccolith at growth Stage 1a. (**b**) Rhombohedral units make up the protococcolith ring, with arrows indicating small V-units present between larger R-units. (**c**) SEM image shows that crystal units appear to be connected by a layer of ribbon-like material, indicated by arrows. (**d**) cryoTEM image of a coccolith at growth Stage 1b. (**e-f**) SEM images show that crystals develop such that they interlock with adjacent units, visible when the coccolith is viewed from below (**e**) and above (**f**). Black arrows point towards the outward facing side of the units. Outer tube and inner tube elements are highlighted by red and blue markers respectively, with the bottom crystal face represented by yellow markers. Arrows indicate the respective direction of growth of each element. (**g**) cryoTEM image of a coccolith at growth Stage 2a. (**h**) Outer and inner tube elements develop, with arrows indicating the growth direction of crystals which causes them to overgrow the adjacent crystal. (**i**) The separation between the outer and inner tube, indicated by yellow arrows, can also be visualized by SEM. Scale bars: 200 nm.

From another undeveloped coccolith, crystal units appear to interlock (Fig. 4d-e), possibly relating to the thickening of spherical mineral related in stage 1b. Crystal growth appears to be primarily in a radial direction, given that an individual unit has its horizontal bottom face outgrow the adjacent crystal in one direction (yellow arrow, Fig. **4e**), while its outer face outgrows the unit on the other side (red arrow, Fig. **4e**). When viewed from above, two separate vertical facets can be observed (red and blue points, Fig. **4f**), with the horizontal bottom face underneath (yellow point, Fig. **4f**). In a coccolith where the tube element has developed at growth Stage 2a (Fig. **4g-h**), two distinct crystal layers become visible (yellow arrows indicating separation, Fig. **4i**), forming the inner and outer tube elements respectively. This two-layered tube element is also a distinguishing feature of *G. oceanica* coccoliths, functioning to tightly interlock crystals around the structure. It is unclear at what point in coccolith development the feature develops however, although it appears in *G. huxleyi* it is in Stage 1 directly after crystals nucleate.

Central area elements were also first observed during the formation of the tube element in Stage 2a (Fig. **5a-d**). CryoTEM and SEM reveals inconsistent morphology in these elements around the ring; either having wider rhombic shapes (red arrow, Fig. **5b**) or being thinner and hammer-shaped (yellow arrow, Fig. **5b**). As central area elements extend inwards in Stage 2b, proximal shield elements first appear (arrows Fig. 5e), and later once adjacent elements start to come into contact in Stage 3a (Fig. **5f-g**) the distal shield elements begin to form (red arrows Fig. 5f). This is in close agreement with the development stages characterised by mapping coccolith morphology using PXCT. Additionally, spindle-like central area elements actually extend from the middle of the inner tube crystals, rather than the base (arrows in Fig. **5h**). This is also concurrent with the position where the central region connects to the coccolith tube in the cryo-PXCT cross-sections (Fig. **2**).

**Figure 5:**
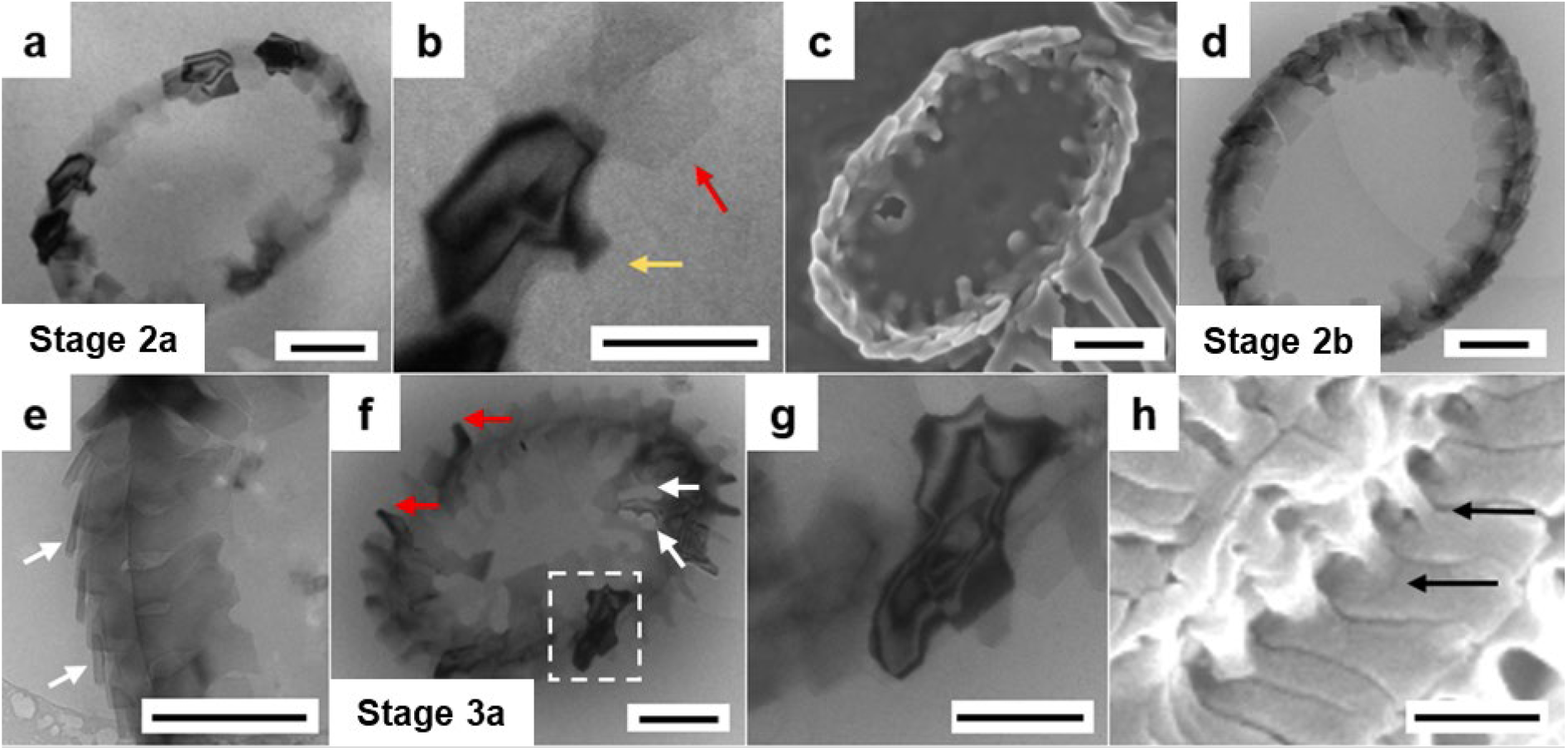
Growth of the central area elements and first appearance of the shield elements across Stages 2a to 3a. (**a**) cryoTEM image of a coccolith at growth Stage 2a. (**b**) Yellow and red arrows highlight central area elements with different morphologies, hammer and rhombic shaped respectively. (**c**) SEM image of a coccolith at growth Stage 2a. (**d**) cryoTEM image of a coccolith at growth Stage 2b. **(e)** Higher magnification of a section of the forming coccolith in Stage 2b, highlighting the inward extension of central area elements and the start of the proximal shield element (white arrows). (**f**) cryoTEM image of a coccolith at growth Stage 3a, where distal and proximal shield elements begin to extend outwards at this stage (**g**). White arrows show gaps between the central elements and red arrows indicate the growing distal shield elements. (**h**) SEM image of a mature coccolith shows spindle-like central area elements (black arrows) extend from the centre of the inner tube crystals. Scale bars: 200 nm.

Stages 3 and 4 of coccolith development concerns the formation of distal and proximal shield elements (Fig. **6a-d**). Radial growth occurs from the upper and lower side of the outer tube element (Fig. **6a-b**); at the bottom to form the plate-like connected crystals of the lower proximal shield (Fig. 6e), and at the top to form the thin spindles that eventually broaden into a T-shape (Fig. **6f-g**). It is challenging to observe crystal morphology in extracted coccoliths at later growth stages due to units being tightly interlocked, with the exact shape of shield elements being revealed when malformations occur (Fig. 6e) or coccoliths break apart during SEM preparation (Fig. 6f-g). In the lower proximal shield, some crystal units have triangular morphology compared to the traditional rectangular shapes. This appears to develop as the crystals grow radially outwards, with some extending faster into the unoccupied space which blocks the space for adjacent shield crystals to grow into (circle, Fig. 6e). The development of anisotropic crystallographic facets as a result of units being tightly interlocked and growing in close contact is commonly observed in coccoliths from multiple species. A similar phenomenon is observed where the upper distal shield elements extend from the outer tube, where growth takes place from half the outer tube crystal (Fig. 6f) as the other section is blocked by the adjacent crystal unit on one side (Fig. 6g). Distal shield elements have crystallographic facets which are not covered by adjacent mineral (Fig. **6g**), meaning confinement does not influence their entire growth. Control over crystal growth in these elements has been strongly linked to coccolith-associated polysaccharides attaching to specific facets and inhibiting further mineral addition, which may be the dominate regulatory mechanism in this region [20-22].

**Figure 6:**
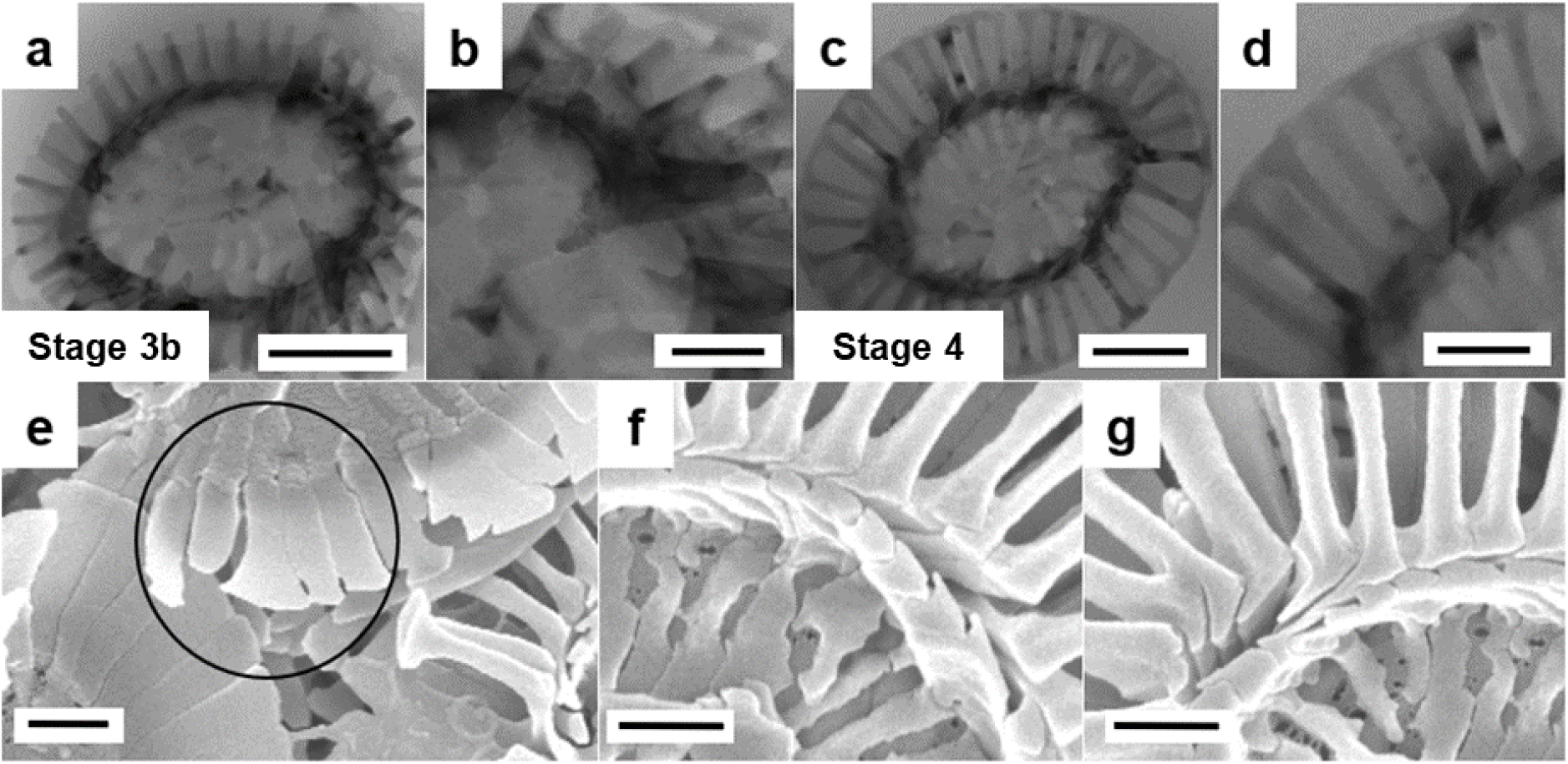
Growth of the distal and proximal shield elements across Stages 3b and 4. (**a-b**) cryoTEM images of a coccolith at growth Stage 3b. (**c-d**) cryoTEM images of a mature coccolith at growth Stage 4. (**e-g**) SEM images of mature coccoliths highlight the morphology of the (**e**) proximal shield (black circle), (**f**) inner tube, and (**g**) distal shield elements. Scale bars: 200 nm.

## 4. Discussion

Despite extensive studies of coccolith growth and its description by the widely used V/R model [14-16], key gaps remain in our understanding of the mechanisms that govern crystal growth. Resolving these requires a technique capable of imaging intracellular mineral intact, across all stages of mineralisation and at sufficient throughput to capture population-level trends. PXCT meets these requirements by enabling three-dimensional imaging of intracellular CaCO_3_ biominerals in their native cellular environment without disrupting the cell [14,15], while providing a large field of view that allows ∼80 cells to be imaged simultaneously. This allowed us to image large numbers of coccoliths and track morphological evolution throughout the entire mineralisation process.

Coccolith morphology largely followed the V/R model, including the exact timings that each element starts and stops growing. We observed that growth roughly occurred in a single direction at one time, starting radial after nucleation before switching to vertical to form the tube element, and finally reverting to radial to produce the shield and central area elements. This suggests that growth direction is controlled by the cell, possibly a consequence of mineralisation being confined to an intracellular vesicle. Confining crystallisation to a specific location is known to be an efficient way to control crystal morphology [42]. Biomineralising organisms can achieve this by precipitating an organic matrix prior to nucleation, while coccolithophores secrete mineral scales in a Golgi-derived specialised vesicle [18]. It is known that specific crystallographic facets and overall crystal morphology in coccoliths can be influenced by the membrane of the vesicle [26, 43], while concentrated gradients created by localised ion channels have been proposed to cause preferential growth of specific facets in this and other species [25, 43].

In addition to providing morphological information, we could infer from the PXCT data that the mass of mature coccoliths range between ∼2 to 3.6 pg. This implies that the cells must mobilize *ca*. 0.8 to 1.4 pg of calcium ions to produce one coccolith. Considering that it takes approximately 60 min for the synthesis of one coccolith, we calculate the uptake and transport of 18-30 fmol calcium ions per hour. These values are in the range of those calculated by Holtz, L.M. et al. [44] and correspond to a flux of ∼5 x 10^6^ calcium ions per min, as previously suggested [45]. It is interesting, though, that half of the coccoliths we analysed were in stage 1, suggesting that the isotropic growth of the calcite crystals in the proto-coccolith ring is the rate-limiting step. Whether this arises from lower calcium ion flux in stage 1 compared to later stages, slower growth kinetics of the crystals during their isotropic growth, the nucleation rate being a limiting factor or other factors, remains to be investigated.

It is plausible that by controlling the shape of the coccolith vesicle, crystal growth can be directed along a specific axis at one time. cryoPXCT revealed that the location of the developing coccolith changes during its formation, suggesting the position of the mineralisation site is under tight cellular control. Although it could not be resolved in the cryoPXCT tomograms, cytoskeletal elements have been proposed to regulate the shape of the coccolith vesicle during mineralisation, therefore it is possible they could dictate the position of the vesicle as well [46, 47]. Further evidence that the location of mineralisation is tightly controlled was also revealed in the consistent positioning of intracellular coccoliths in regions of the cell not covered by the external coccosphere layer. The cytoskeleton has also been suggested to control exocytosis, further strengthening an argument that it could provide additional control of coccolith position to ensure that mature scales exit the cell in an empty space in the external coccosphere. In *Coccolithus braarudii*, coccoliths were shown to be secreted from a singular location, with the cell rotating within the coccosphere to ensure that the scale exited into an empty space [8]. This suggests that the mineralisation site remains at a constant location, with coccosphere assembly influenced more by cell movements and the position of external coccoliths.

Previous coccolith growth studies generally missed the formation of the entire tube element. In the coccoliths produced by related species *G. oceanica*, the outer and inner tube elements are responsible for many structural coccolith features, growing at tilted vertical angles in opposite directions causing adjacent crystal units to overlap and tightly interlock around the ring [23]. These features were not included in the original V/R model despite their now apparent importance to overall crystal morphology and organisation. While we cannot confirm the outer and inner tube have the same morphology in *G. huxleyi*, an assumption can be made that they form in a similar way given the comparable structures of the coccoliths produced by both species. Additionally, both the tube layers have similar morphologies in each species, the inner crystals rhombohedral and the outer plate-like, while we also observed tightly interlocked crystals at early stages of the tube element formation in *G. huxleyi* coccoliths. Similar 3D reconstruction and analysis of *G. huxleyi* coccolith structure should be conducted to confirm this configuration of crystal elements.

Interlocking crystals are a feature of coccolithophore biominerals, equally observed in *C. carterae* [19, 24], *C. leptoporus* [25] and *G. oceanica* [23]. In *G. huxleyi*, as shown here the first stages of crystal growth create a ring of interlocking units, which could be a mechanism that is generic across all coccolithophores species. In the coccoliths produced by *C. carterae*, constrained growth spaces are known to dictate the overall morphology of crystal units [24]. During the early stages of *G. huxleyi* formation, we observe rhombohedral units growing laterally towards each other, with the confined space created by the adjacent unit blocking certain regions and influencing where mineral can be added. This may originate from crystals nucleating at slightly offset angles to enable their organisation in a ring [17, 24, 25], possibly causing adjacent units to block others which prevents mineral from expanding into certain regions. It is likely that control over growth begins directly after the crystals nucleate, as the tube element in *G. oceanica* separates into its two layers around this stage. The influence of adjacent units affecting growth space and crystal morphology in *G. huxleyi* is also evident in central area and shield elements. Growth of these elements appears to occur at faster rates in some crystal units, causing them to outgrow others and block their expansion, changing the morphology of the mineral that forms as a result. Distal shield elements also have their morphology partially controlled by confined growth space, with their formation starting from the regions on the outer tube crystals that are not blocked by adjacent units.

Organic macromolecules are also known to have a significant influence on crystal growth and morphology, although it is difficult from this study to quantify the extent to which macromolecular occlusion might influence different stages of *G. huxleyi* coccolith growth. Polysaccharides have been found occluded in *G. huxleyi* coccoliths, located either in the crystal lattice or at the grain boundaries [22, 48]. *In vitro* experiments have demonstrated that these macromolecules favour binding to acute calcite edges to promote crystal elongation along the crystallographic c-axis [20, 21]. As promoted growth is radial to the base plate rather than vertical, organic control is likely a greater factor in the formation of crystal elements in these directions, such as the growth of central elements and the proximal and distal shields. Further information is required on location of organic structures within the coccolith crystals to determine if and how they control mineral growth.

Coccolithophores are not just important organisms to study for their design of functional materials, but also have important applications as geological proxies for both palaeoclimate reconstructions and models for future climates. For coccolithophores to be a good proxy, a thorough understanding of coccolith mineralisation is required, including knowledge of the cellular mechanisms involved in growth as well as accurate information about mineral composition and weight. Here, we provide an improved estimation of individual coccolith volume and weight at different stages of formation, which could be a valuable resource to ecological studies calculating carbon isotope ratios or determining the influence of increased CO_2_ seawater concentration on coccolithophore calcification.

## 5. Conclusions

We have provided new findings in the biomineral formation process of the coccolithophore *G. huxleyi*. The length-scale overarching combination of cryo-PXCT and electron microscopy allowed us to follow the formation of calcite coccolith scales in their native environment in 3D over their entire mineralisation cycle. The crystal subunits assemble into a tightly confined spherical ring, the confinement of this geometric arrangement strongly influences their growth, leading to the development of crystallographic facets that reflect the presence of neighbouring subunits. Additional control appears to be imparted by controlling the position of the mineralisation site in the cell, currently suggested to function as a mechanism to stop vertical growth, while also ensuring that mature coccoliths are exocytosed in the correct position to form the coccosphere. This study highlights the high degree of cellular control imparted during mineralisation, often employing pre-existing cellular pathways to ensure that crystalline materials display the functional properties required by organisms to survive in their environment.

## Acknowledgements

Funding for this research was provided by NERC through an E4 DTP studentship (NE/S007407/1) to F.N. and by the European Research Council (European Union’s Horizon H2020 research and innovation program grant agreements No 724881 to V.C and T.G. CryoPXCT was performed at the coherent small-angle x-ray scattering (cSAXS) beamline at the Swiss Light Source at the Paul Scherrer Institut, Proposal 20200711. SEM data acquisition and cryoPXCT tomogram segmentation using Avizo were performed in the cryoFIB-SEM facility at the University of Edinburgh (EPSRC grant No, EP/P030564/1). CryoTEM data acquisition was supported by funding for the Wellcome Discovery Research Platform for Hidden Cell Biology [226791s] and we gratefully acknowledge support from the Structural Biology core.

## Notes

### Competing Interest Statement

The authors have declared no competing interest.

