## Supplementary data for "Deciphering Coccolith Formation: Advanced Microscopy Insights from the Biomineralisation of *Gephyrocapsa huxleyi*"

**This PDF file includes:**

Tables S1 and S2

Figs. S1 to S5

Tables S1 to S2

Legend for Movie S1

**Other Supplementary Materials for this manuscript include Movie S1.**

**Supporting Table 1**. Scan parameters used for the CryoPXCT dataset

| Parameter | 456 |
| --- | --- |
| FOV [µm^2^] | 56 x 15 |
| Number of Angles | 1400 |
| Step size [µm] | 1.2 |
| Exp time [s] | 0.025 |
| Dose [Gy] | 4.33 x 10^7^ |
| FSC Resolution [nm] | 56.3 |

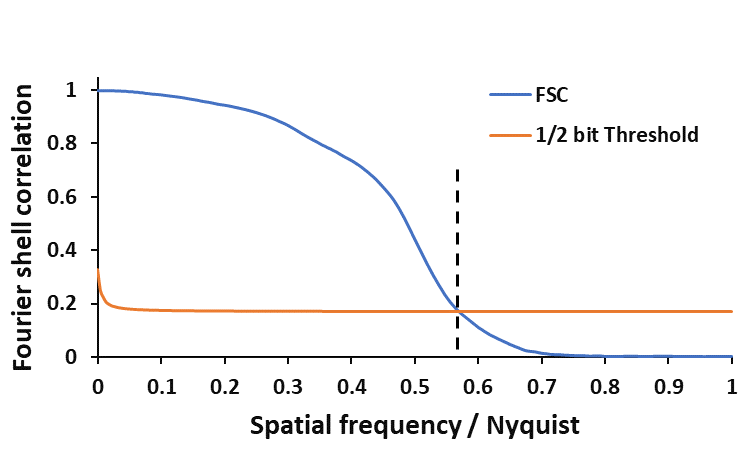

**Supporting Figure 1**. **Spatial-Resolution of Ptychographic Tomograms obtained from the *G. huxleyi* dataset.** Fourier shell correlation (FSC) line plots of the acquired electron density tomograms. This estimation was performed on a region of the sample that excludes the capillary contribution. The selected threshold for determining the resolution is the ½ bit criterion. The voxel size for the tomogram is 32 nm. The half-period spatial resolution estimate for all samples is ~56 nm.

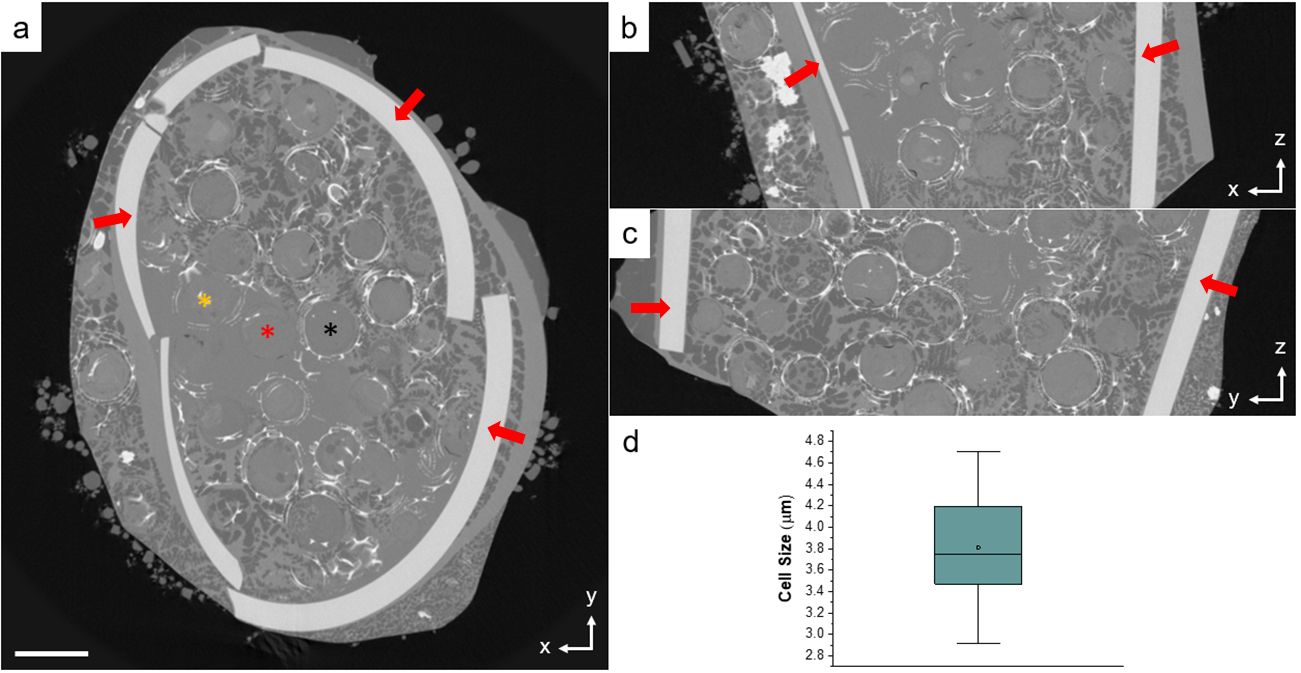

**Supporting Figure 2: CryoPXCT imaging of *G. huxleyi* cells.** CryoPXCT tomogram of a glass capillary containing *G. huxleyi* cells, shown across the *xy* (**a**), *xz* (**b**) and *yz* (**c**) planes. Inside the capillary, *G. huxleyi* cells were actively calcifying, indicated by the presence of a nearly full (black marker) or partial (yellow marker) coccosphere, while some cells displayed no external coccoliths (red marker). The red arrows in (**a-c**) indicate the wall of the capillary. (**d**) A box-and-whisker plot represents the size of all cells in the capillary. Scale bar is 5 µm.

**Movie S1**

The full cryoPXCT tomogram and 3D rendering of the glass capillary containing *G. huxleyi* cells is shown in **Movie S1**. For clarity, we display the 3D rendering of only a limited number of extracellular mature and intracellular forming coccoliths that are representative of the sample. The coccoliths were segmented using the following colour code:

Stage 1: pink (stage 1a) and light blue (stage 1b)

Stages 2a and 2b: olive green

Stages 3a and 3b: dark green

Stage 4 (mature, intracellular): dark blue

Stage 5 (extracellular): yellow

**Supporting Table 2: Intracellular coccolith stages.** The number of intracellular coccoliths at each stage of formation across the population of *G. huxleyi* cells, as revealed by cryo-PXCT. Growth stages are defined by the appearance of certain elements and overall morphology, as defined in Fig. 2.

| Stage | Number of coccoliths at each stage |
| --- | --- |
| 1 | 45 |
| 2 | 11 |
| 3 | 17 |
| 4 | 15 |

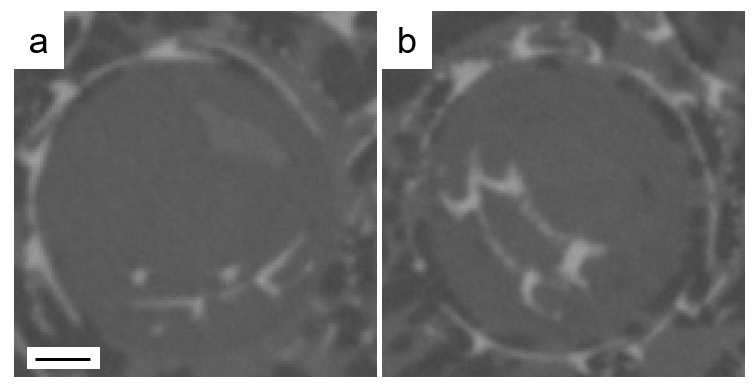

**Supporting Figure 3: Synthesis of multiple coccoliths within a single *G. huxleyi* cell.** Cross-sections through separate *G. huxleyi* cells show the presence of multiple intracellular coccoliths. When this observed, one coccoliths was fully mature, while the other was either at the protococcolith stage (a), or more developed (b). Scale bar is 0.5 µm.

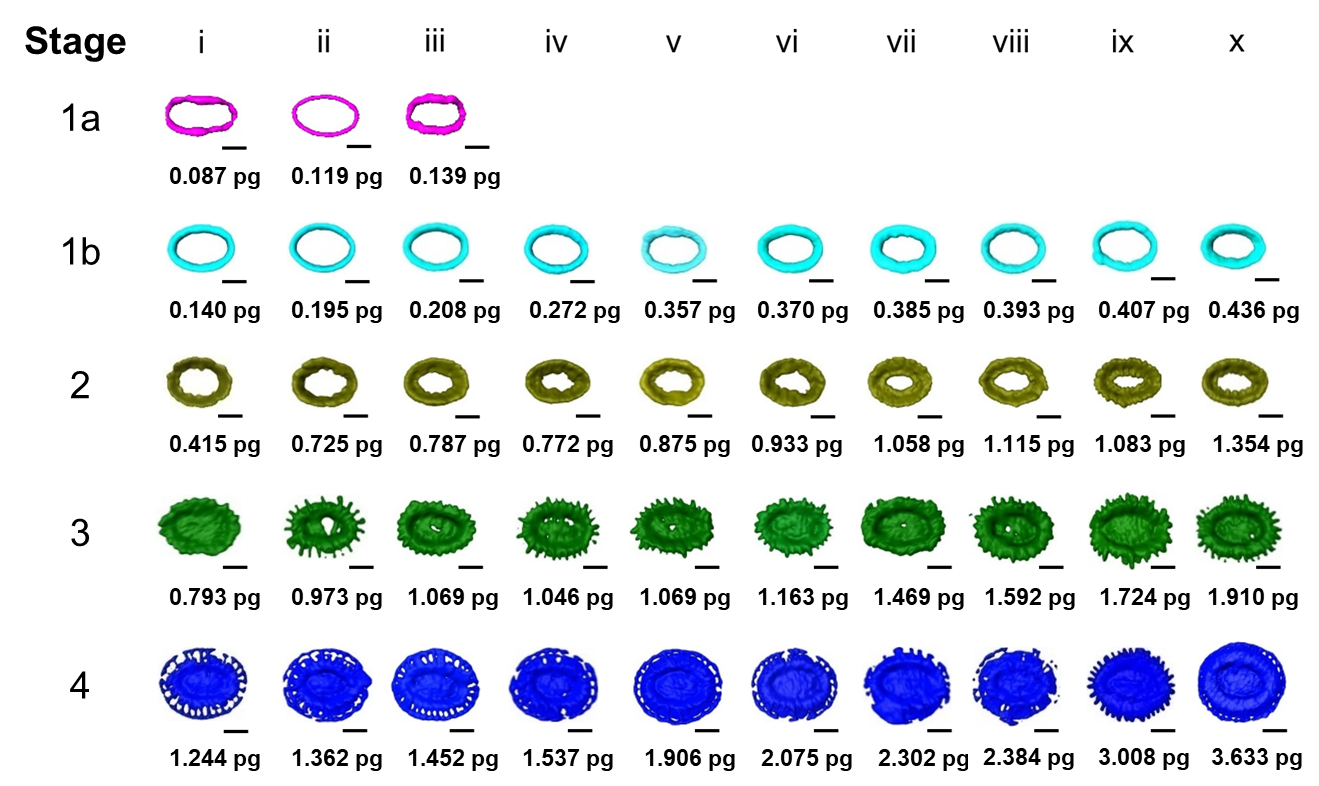

**Supporting Figure 4: Mineral mass of coccoliths.** Volume renderings of all intracellular coccoliths analysed using PXCT for each growth stage. The mass of each coccolith is shown below. Scale bars are 500 nm.

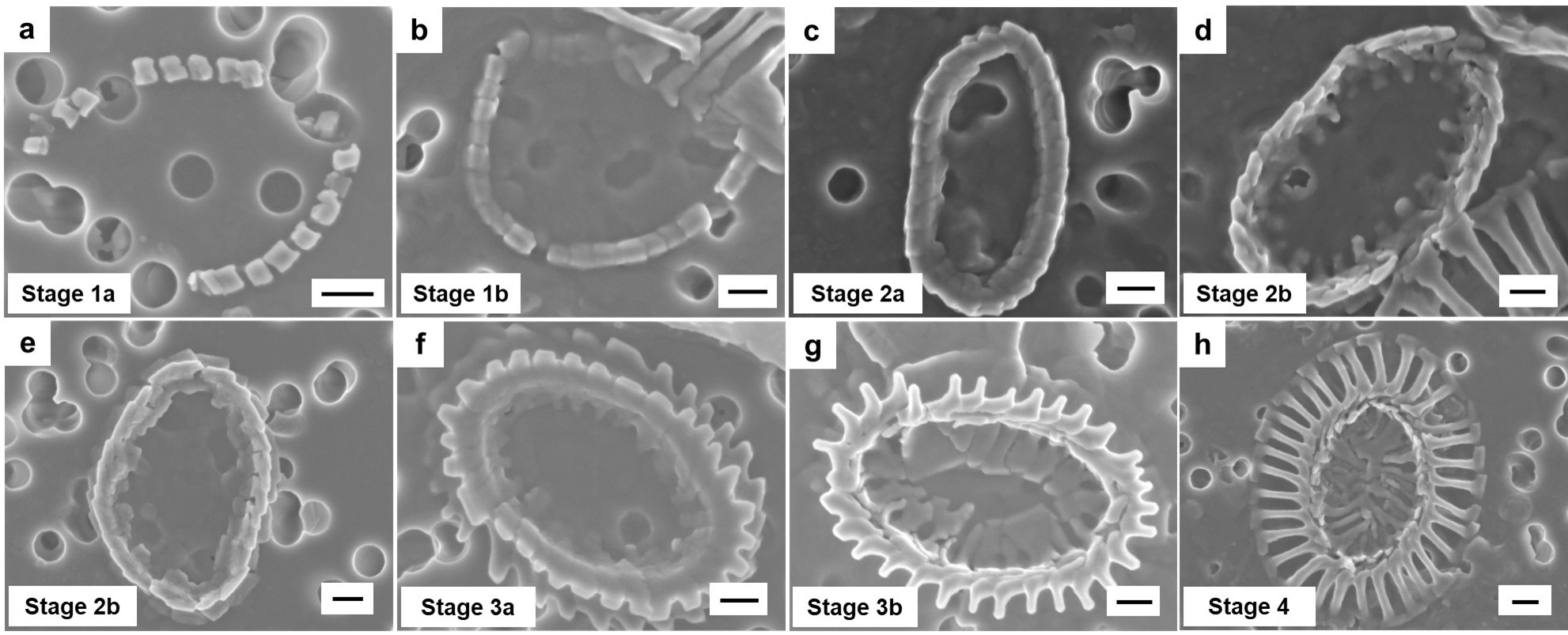

**Supporting Figure 5: Coccolith crystal growth in *G. huxleyi*:** Intracellular *G. huxleyi* coccoliths at various stages of development were imaged by scanning electron microscopy (SEM). Growing coccoliths were extracted from within cells by osmotic pressure. (a) At the earliest stage of mineralization, small crystals nucleate around the ring in stage 1a. (b) These expand to form a continuous ringed structure in stage 1b. (c) Units become interlocked and extend vertically in stage 2a, before growth of inner elements starts in stage 2b (d). (e) The inner regions extend further inward, before outer elements, first the proximal shield in stage 2b, then the distal shield in stage 3a (f), become visible. (g) Further growth of these elements can be observed in stage 3b, before full extension results in the formation of mature coccolith scales in stage 4 (h). Growth of the crystal units follows what is observed from ptychographic tomography and cryoTEM.
